# Efficacy of GC-376 against SARS-CoV-2 virus infection in the K18 hACE2 transgenic mouse model

**DOI:** 10.1101/2021.01.27.428428

**Authors:** C. Joaquín Cáceres, Stivalis Cardenas-Garcia, Silvia Carnaccini, Brittany Seibert, Daniela S. Rajao, Jun Wang, Daniel R. Perez

**Author notes:** Co-corresponding authors: Department of Pharmacology and Toxicology, College of Pharmacy, University of Arizona. 1703 E. Mabel St, Tucson, AZ 85721., Department of Population Health, Poultry Diagnostic and Research Center, University of Georgia. 953 College Station Road, Athens, GA 30602.

## Abstract

The COVID-19 pandemic caused by the Severe Acute Respiratory Syndrome Coronavirus-2 (SARS-CoV-2) is the defining global health emergency of this century. GC-376 is a M^pro^ inhibitor with antiviral activity against SARS-CoV-2 *in vitro*. Using the K18-hACE2 mouse model, the *in vivo* antiviral efficacy of GC-376 against SARS-CoV-2 was evaluated. GC-376 treatment was not toxic in K18-hACE2 mice and produced milder tissue lesions, reduced viral loads, fewer presence of viral antigen, and reduced inflammation in comparison to vehicle-treated controls, most notably in the brain in mice challenged with a low virus dose. Although GC-376 was not sufficient to improve neither clinical symptoms nor survival, it did show a positive effect against SARS-CoV-2 *in vivo*. This study supports the notion that the K18-hACE2 mouse model is suitable to study antiviral therapies against SARS-CoV-2, and GC-376 represents a promising lead candidate for further development to treat SARS-CoV-2 infection.

## Introduction

The coronavirus disease (COVID-19) pandemic caused by the Severe Acute Respiratory Syndrome Coronavirus-2 (SARS-CoV-2) is the defining global health crisis of our time reaching over 45 million cases worldwide by October 2020 ^1^. SARS-CoV-2 belongs to the Coronaviruses family, which are RNA viruses commonly associated with mild upper respiratory illness. In the past few years, novel coronaviruses have emerged from animal reservoirs crossing the species barrier causing sporadic outbreaks in humans such as SARS-CoV and the Middle Eastern Respiratory Syndrome (MERS). The current global health emergency prompted a race to develop resources to combat the pandemic. Several vaccines against SARS-CoV-2 are currently in different stages of pre-clinical or clinical evaluation ^2-4^. Among these, the Pfizer/BioNTech’s mRNA-based vaccine against SARS-CoV-2 has been approved for human use in the United Kingdom and awaits approval in the U.S. and other countries. Additionally, a number of viral proteins and host factors have been proposed as small molecule antiviral drug targets to combat SARS-CoV-2 infection. Among those, the viral main protease (M^pro^) is one of the most extensive explored drug targets^5^, and a number of M^pro^ inhibitors are in different stages of preclinical and clinical development. GC-376 is a representative M^pro^ inhibitor that has shown antiviral activity against FIP CoV in experimentally infected cats ^6,7^. The ∼30kb viral genome SARS-CoV-2 genome encodes two polyproteins, pp1a and pp1ab, which must be cleaved in order to be functional and lead to active viral replication. Cleavage is mediated by the M^pro^ and the papain-like (PL^pro^) proteases ^8^, both potential antiviral targets. *In vitro* studies have shown inhibition of SARS-CoV M^pro^ in the presence of GC376 ^9^. More recently, the potential antiviral activity of GC-376 against SARS-CoV-2 has been demonstrated *in vitro* placing GC-376 as a promising antiviral candidate for treatment of SARS-CoV-2 infections ^10-13^. However, the antiviral activity of GC-376 against SARS-CoV-2 *in vivo* has not been addressed, neither have any other M^pro^ inhibitors that are currently in development. Therefore, to fulfil the knowledge gap and provide insights of the translational potential of M^pro^ inhibitors as SARS-CoV-2 antivirals, we report in this study the establishment of the SARS-CoV-2 infection mouse model and the *in vivo* antiviral efficacy of GC-376 using this model. A roadblock for *in vivo* studies involving SARS-CoV-2 has been the limited information regarding suitable animal models, although research is clearly moving fast in this particular area. SARS-CoV-2 is able to infect different animal models including hamsters, ferrets, and non-human primates. However, the severity of the disease in these models only ranges from mild to moderate, which makes it harder for the assessment of efficacy of antivirals or vaccines ^14-17^. A similar problem was experienced for the study of SARS-CoV-1, which was in part resolved by the development of a lethal mouse model of infection for SARS-CoV-1 by introducing the human angiotensin-converting enzyme type 2 (hACE2) receptor under the control of the keratin 18 promoter (K18-hACE2). Using this model, high mortality was observed upon SARS-CoV-1 infection ^18-20^. Since SARS-CoV-2 also utilizes the hACE2 receptor, the K18-hACE2 has been suggested as a suitable mouse model for the study of SARS-CoV-2 and recent reports show the development of clinical signs and mortality after SARS-CoV-2 challenge ^20,21^. Thus, the K18-hACE2 mouse model was chosen to evaluate the antiviral efficacy of GC-376 upon SARS-CoV-2 infection. Two different virus challenge doses of SARS-CoV-2 were used showing clinical signs of disease and mortality that correlated with the dose administered. Modest differences in terms of clinical signs, weight loss, and survival were observed between GC-376 and vehicle mice. However, further analyses indicated that GC-376/SARS-CoV-2 challenged mice showed lower viral loads, milder tissue lesions and reduced inflammation compared to vehicle SARS-CoV-2 challenged controls showing that GC-376 can act as a SARS-CoV-2 antiviral *in vivo*.

## MATERIALS AND METHODS

### Ethics Statement

Studies were approved by the Institutional Animal Care and Use Committee. Studies were conducted under animal biosafety level 3 containment. Animal studies and procedures were performed according to the Institutional Animal Care and Use Committee Guidebook of the Office of Laboratory Animal Welfare and PHS policy on Humane Care and of Use of Laboratory Animals. Animal studies study were carried out in compliance with the ARRIVE guidelines (https://arriveguidelines.org). Animals were humanely euthanized following guidelines approved by the American Veterinary Medical Association (AVMA).

### Cells and Virus

Vero E6 Pasteur were kindly provided by Maria Pinto (Center for Virus research, University of Glasgow, Scotland, UK). Cells were maintained in Dulbecco’s Modified Eagles Medium (DMEM, Sigma-Aldrich, St Louis, MO) containing 10% fetal bovine serum (FBS, Sigma-Aldrich, St Louis, MO), 1% antibiotic/antimycotic (AB, Sigma-Aldrich, St Louis, MO) and 1% L-Glutamine (Sigma-Aldrich, St Louis, MO). Cells were cultured at 37°C under 5% CO_2_. SARS-CoV-2 (Isolate USA-WA1/2020) was kindly provided by Dr. S. Mark Tompkins, Department of Infectious Diseases, University of Georgia. Viral stocks were prepared in Vero E6 Pasteur cells. Briefly, cells in a T75 flask were incubated 1 h with 1 ml of viral inoculum. After incubation, the inoculum was removed and cells were cultured with DMEM containing 2% FBS, 1% AB. After 96 h, the supernatant was collected, centrifuged at 15,000 g for 15 min., aliquoted and stored at -80°C until use. Viral stocks were titrated by tissue culture infectious dose 50 (TCID_50_) and virus titers were established by the Reed and Muench method ^22^.

### Chemical compounds

GC-376 (CAS: 1416992-39-6) was purchased from Enovation chemicals (Green book, NY) and dissolved in deionized water (HyClone, VWR, West Chester, PA) to the desired concentration.

### Mouse studies

6-week-old female K18-hACE2 mice were purchased from Jackson laboratories (Bar Harbor, ME). Mice were randomly distributed in the number of groups (Fig 1A), anesthetized with isoflurane and challenged intranasally (i.n.) with 50 µl of either phosphate buffer saline (PBS), or 1×10^3^ TCID50/mouse or 1×10^5^ TCID50/mouse. At 3 h post-challenge, GC-376 (20 mg/kg/dose, 40 mg/kg daily) or vehicle (H_2_O) was administered to each mouse through intraperitoneal injection (i.p.) and continued for 7 days, twice per day. Mice were monitored twice daily for clinical signs of disease along the entire course of the experiment. Mice that lost ≥25% of their initial body weight (a score of 3 on a 3-point scale of disease severity) were humanely euthanized. At 2 and 5 dpc, subsets mice (n=2/time point from G1 and G2 and n=3/time point from G3, G4, G5, and G6) received an overdose of isoflurane, and were humanely euthanized. Lungs, nasal turbinates (NT), spleen, liver, small intestine (SI) and brain were collected from each mouse post-mortem. Tissues were stored at -80°C until further analysis. At 14 dpc, the same procedure was performed with all the remaining animals.

**Figure 1.**
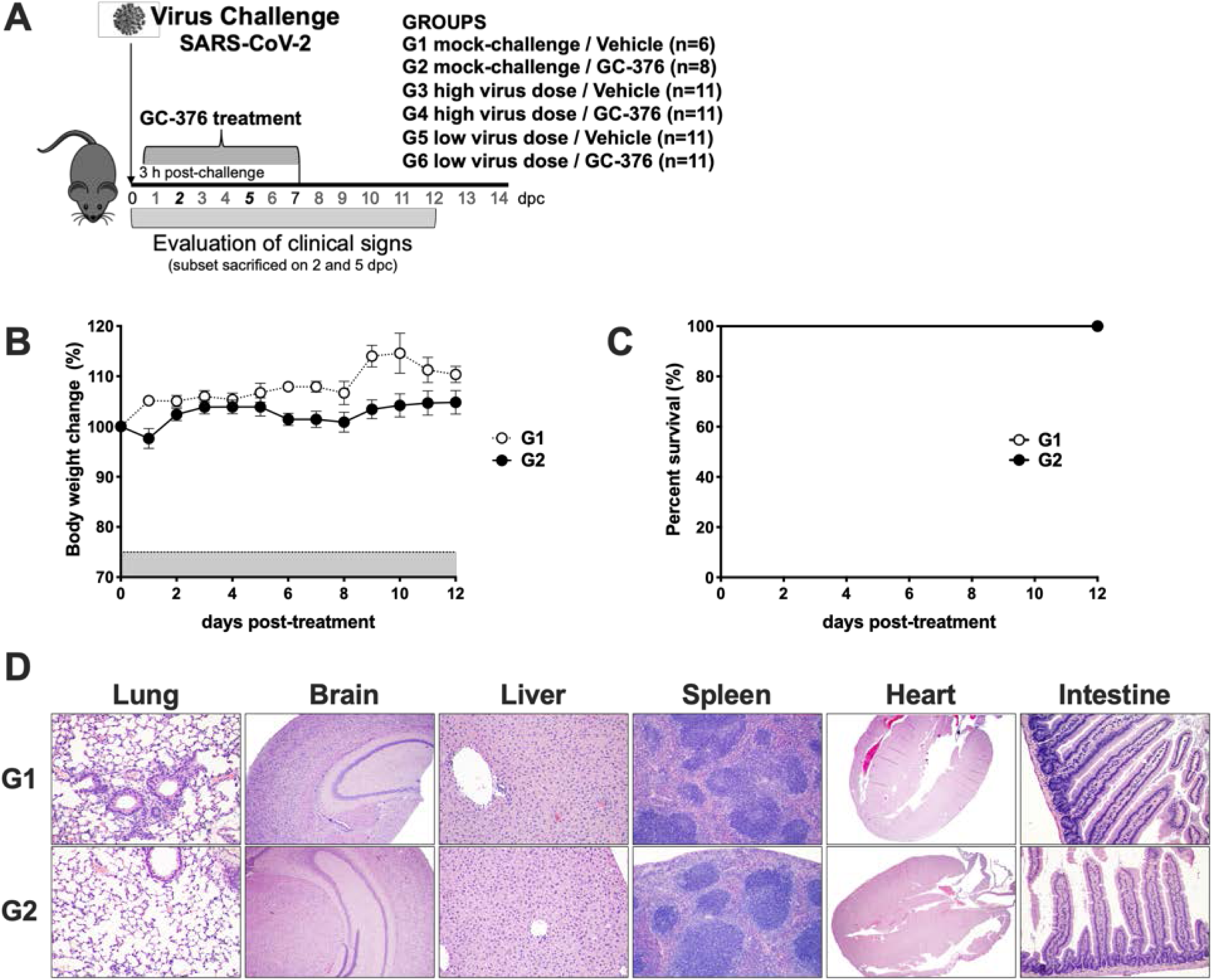
Schematic representation of the experimental design and evaluation of the safety of GC-376 in K18-hACE2 mice. **(A)** K18-hACE2 were either mock challenged (G1 and G2) or challenged with SARS-CoV-2 at either a high virus dose (1×10^⋀^5 TCID50/mouse; G3 and G4) or a low virus dose (1×10^⋀^3 TCID50/mouse; G5 and G6). I.P. treatment with vehicle (G1, G3, and G5) or GC-376 (G2, G4, and G6, 40mg/kg day). Subsets of mice were humanely euthanized at 2 and 5 dpc and different tissues were collected. Mice (n=8) received 40mg/kg of GC-376 split in two doses per day for 7 days and **(B)** weight changes and **(C)** survival were monitored for 12 days. **(D)** Mice were humanely euthanized at 14 dpc and lungs, brain, liver, spleen, heart and SI were collected. HE slides were generated and analyzed in comparison with the negative control group. Representative pictures were taken at 20X magnification for all tissues except brain samples that were taken at 4X.

### Tissue sample preparations

Tissues homogenates were generated using the Tissue Lyzer II (Qiagen, Gaithersburg, MD). Briefly, 500 µl of PBS-AB was added to each sample along with Tungsten carbide 3 mm beads (Qiagen). Samples were homogenized during 10 min and then centrifuged at 15,000 g for 10 min. Supernatants were collected, aliquoted and stored at -80°C until further analysis.

### RNA extraction and RT-qPCR

RNAs were extracted from the different tissue homogenates using the MagMax-96 AI/ND viral RNA isolation kit (ThermoFisher Scientific, Waltham, MA) following manufacturer’s protocol. A one-step real time quantitative PCR (RT-qPCR) based on the Nucleoprotein gene segment was used as surrogate of viral load and it was employed using the primers 2019-nCov_N2-F (5’-TTACAAACATTGGCCGCAAA-3’) and 2019-nCov_N2-R (5’-GCGCGACATTCCGAAGAA-3’). A probe with FAM as a reporter and TAMRA as a quencher was used (5’-FAM-ACAATTTGCCCCCAGCGCTTCAG-TAMRA-3’). The RT-qPCR was performed in a QuantStudio 3 Real-Time PCR System (ThermoFisher Scientific, Waltham, MA) using a Quantabio qScriptTM XLT One-Step RT-qPCR ToughMix kit (Quantabio, Beverly, MA) in a final reaction volume of 20 μl. Each reaction mixture contained 1X master mix, 0.5 μM forward and reverse primers, 0.3 μM probe, and 5 μl of RNA. The qPCR cycling conditions were 50°C, 20 min; 95°C, 1 min, 40 cycles at 95°C, 1 min; 60°C, 1 min; and 72°C 1 s; with a final cooling step at 4°C C. A standard curve was generated using 10-fold serial dilutions of a SARS-CoV-2 virus stock of known titer to correlate RT-qPCR crossing point (Cp) values with the viral load from each tissue. Viral loads were calculated as Log_10_ TCID50 equivalents/per gram of tissue.

### Histopathology and immunohistochemistry

Selected tissues, including lungs, NT, spleen, liver, heart, SI, and brain were collected from a representative number of mice in each group and at different timepoints (2, 5 and 14 dpc) for histopathological examination. Tissues were placed in 10% neutral-buffered formalin (NBF), fixed for at least 72 hours, paraffin embedded and processed for routine histopathology with hematoxylin and eosin staining (HE). Tissues were subjectively scored by a pathologist as none (-), mild (+), mild to moderate (++), moderate (+++), moderate to severe (++++) and severe (+++++) based on lesion severity and extent of inflammation. Additionally, antibodies targeting the SARS-CoV-2 nucleoprotein (N) (ThermoFisher Scientific, Waltham, MA; dilution 1/500), the CD3 receptor (Agilent technologies, Santa Clara, CA; dilution 1/1000) and the allograft inflammatory factor 1 (IBA-1) (Fujifilm WAKO chemicals, Richmond, VA; dilution 1/8000) were also used to perform immunohistochemistry (IHC) on selected tissues. The intensity and distribution of the staining was used to estimate the amount of viral N, CD3 and IBA-1 antigens which was subjectively scored by a pathologist using a scale from none (-) to large amount (+++++), being the large amount the highest level of positivity.

### Graphs/Statistical analyses

All data analyses and graphs were performed using GraphPad Prism software version 8 (GraphPad Software Inc., San Diego, CA). For multiple comparisons, two-way analysis of variance (ANOVA) was performed. A *P* value below 0.05 was considered significant.

## RESULTS

### Modest effect of GC-376 treatment on the clinical outcome of disease in K18-hACE2 mice challenged with SARS-CoV-2

GC-376 has been shown to be effective against FIP CoV in cats ^6^ and against SARS-CoV-2 *in vitro* ^11,12^. We evaluated the efficacy of GC-376 against SARS-CoV-2 *in vivo* using the K18-hACE2 mouse model (Fig 1A). Concomitantly, we evaluated the potential toxicity of GC-376 in K18-hACE2 mice, noting that previous studies have shown no liver toxicity caused by GC-376 in other mouse strains at the dose administered in this study ^23^. Thus, K18-hACE2 mice were either mock-challenged (PBS, G1, n=6/group and G2, n=8/group), challenged with a high dose of SARS-CoV-2 virus (1×10^5^ TCID50/mouse, G3 and G4, n=11/group), or challenged with a low dose of SARS-CoV-2 virus (1×10^3^ TCID50/mouse, G5 and G6, n=11/group). At 3 h post-challenge, mice were treated for 7 days, twice daily i.p. with either vehicle (G1, G3, and G5) or GC-376 (G2, G4, and G6, 40 mg/kg per day, 20 mg/kg each dose). No significant weight loss and 100% survival was observed in both the mock-challenged mice treated with GC-376 (G2) and the mock-challenged vehicle control group (G1, Figs 1B and 1C). Furthermore, lungs, brain, liver, SI, spleen and heart did not present any tissue alteration at 14 days post-treatment with GC-376 in comparison with the vehicle control group (Fig 1D). In mice challenged with the high virus dose (G3 and G4), there was a period of relatively normal activity (response to the environment and stimulus given by the personnel) and physical appearance (Fig 2A and B) followed by rapid weight loss and presentation of clinical signs (Fig 2C). By 6 dpc, mice in G3 (high virus dose/vehicle, white symbols) showed about 20% weight loss and by 8 dpc there were no survivors because they have either succumbed to the infection or had to be humanely euthanized due to severe clinical signs (lethargy, ruffled fur, labored breathing, and/or ≥25% body weight loss, Fig 2D). These observations are consistent with previous reports^20,21,24,25^. G4 mice (high virus dose/GC-376) showed similar clinical signs but such progression was delayed by about 24 h (∼15% weight loss by 6 dpc). In terms of body weight loss, statistically significant differences were established between G3 and G4 from 4 to 6 dpc. Of note, the G4 mice (high virus dose/GC-376) showed 20% survival (n=1 out of 5) but such difference was not statistically significant.

**Figure 2.**
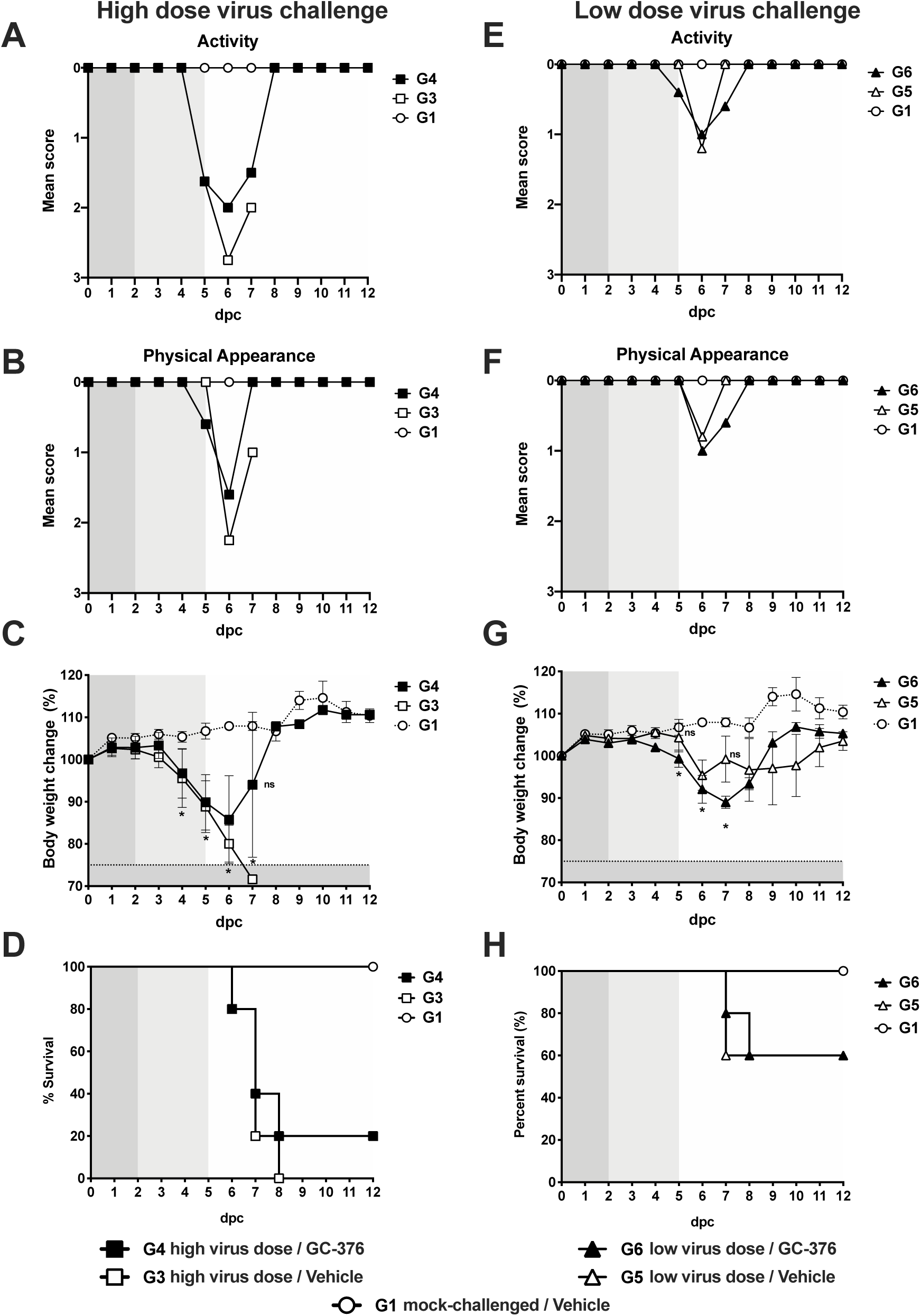
Efficacy of GC-376 in terms of clinical signs and survival after challenge with SARS-CoV-2. Mice (n=11/group) were challenged with a **(A-D)** high dose (1×10^⋀^5 TCID50/mouse) or **(E-H)** low dose (1×10^⋀^3 TCID50/mouse) of SARS-CoV-2 and treated with drug vehicle (white symbols) or 40mg/kg of GC-376 split in two doses per day for 7 days (black symbols). **(A and E)** activity, **(B and F)** physical appearance, **(C-G)** weight change and **(D and H)** survival (n=5). Statistical analysis was performed by a two-way ANOVA, * p<0.05. Grey shading represents remaining number of animals at time of clinical evaluation (n=11; dark grey, n=8; mid grey, n=5 light grey)

We analyzed whether the antiviral activity of GC-376 could improve disease outcome when a low challenge dose of SARS-CoV-2 was used. As expected, progression of the disease occurred slower compared to groups of mice challenged with the high virus dose (G3 and G4, Fig 2A-D). It was interesting to observe transient reduced activity and deterioration of physical appearance (Fig 2E and F) regardless of treatment accompanied by consistent and statistically significant body weight loss between 5 to 7 dpc in all mice regardless of group at the low virus challenge dose compared to mock-challenged/vehicle treated mice, ultimately leading to 60% survival (Fig 2G and H). These observations are significant because it suggests that disease progression due to SARS-CoV-2 infection in the k18-hACE2 mouse model is virus dose dependent adding more value to the system to better study SARS-CoV-2 pathogenesis and host responses. Of note, a trend towards increased weight loss but faster body weight recovery was observed in G6 mice (low virus dose/GC-376) compared to G5 mice (low virus dose/vehicle). However, and despite the 100 times lower virus challenge dose, we did not notice an improvement in the clinical outcome of mice treated with GC-376 compared to mice in vehicle control group. Taken together, these results suggest that GC-376 has modest benefit in K18-hACE2 mice infected with SARS-CoV-2 in terms of clinical signs outcome, weight changes, and survival.

### A trend towards reduced SARS-CoV-2 vRNA loads in GC-376-treated mice, particularly in the brain

To better characterize the effect of GC-376 treatment against SARS-CoV-2, sections of NT, lungs, brain, liver, small intestine (SI) and spleen were collected at 2 and 5 dpc and each tissue split into two to assess vRNA viral loads and for histopathological analyses, respectively. Tissue homogenates were prepared, vRNA extracted and then vRNA loads and vRNA tissue distribution were evaluated by RT-qPCR. High vRNA content consistent with active virus replication was observed in tissue homogenates from the NT, lungs, and brain of mice in G3 (high virus dose/vehicle). RT-qPCR results were consistent with the clinical observations in G3 mice, with average higher vRNA signals at 2 dpc than at 5 dpc in samples from the lungs and NT, but the opposite in other tissues (brain, liver, SI and spleen) indicative of systemic virus spread (Fig 3A). Likewise, vRNA loads in G5 mice (low virus dose/vehicle) followed disease progression with higher average signals on 5 dpc compared to 2 dpc, particularly in the brain where these differences were statistically significant (Fig 3A). Also consistent with the clinical outcome of the disease, reduced vRNA load averages were observed in NT, lungs and brain samples from mice in G4 (high virus dose/GC-376) compared to homologous samples from mice in G3 (high virus dose/vehicle). Samples from mice in G4 showed more individual variations in vRNA loads, several at levels below limit of detection, than the counterparts in G3 (Fig 3B). Similarly, more individual variations in vRNA levels were observed in NT, lungs and brain samples from G6 mice (low virus dose/GC-376) than in G5 mice (low virus dose/vehicle) (Fig 3C). Interestingly, brain samples from G6 mice showed statistically significant lower vRNA levels than those from G5 mice. Overall, these results suggest high levels of SARS-CoV-2 replication in the NT, lungs, and brain of K18-hACE2 mice and detectable vRNA in liver, SI and spleen. GC-376 treatment leads towards reduced virus load averages with the caveat of noticeable individual variations among treated animals, and the inability to properly prevent the burden and mortality caused by SARS-CoV-2 in K18-hACE2 mice. At the low dose virus challenge, GC-376 prevented to great extent the virus’ ability to reach the brain, but such effect was not correlated with increased survival compared to vehicle-treated controls.

**Figure 3.**
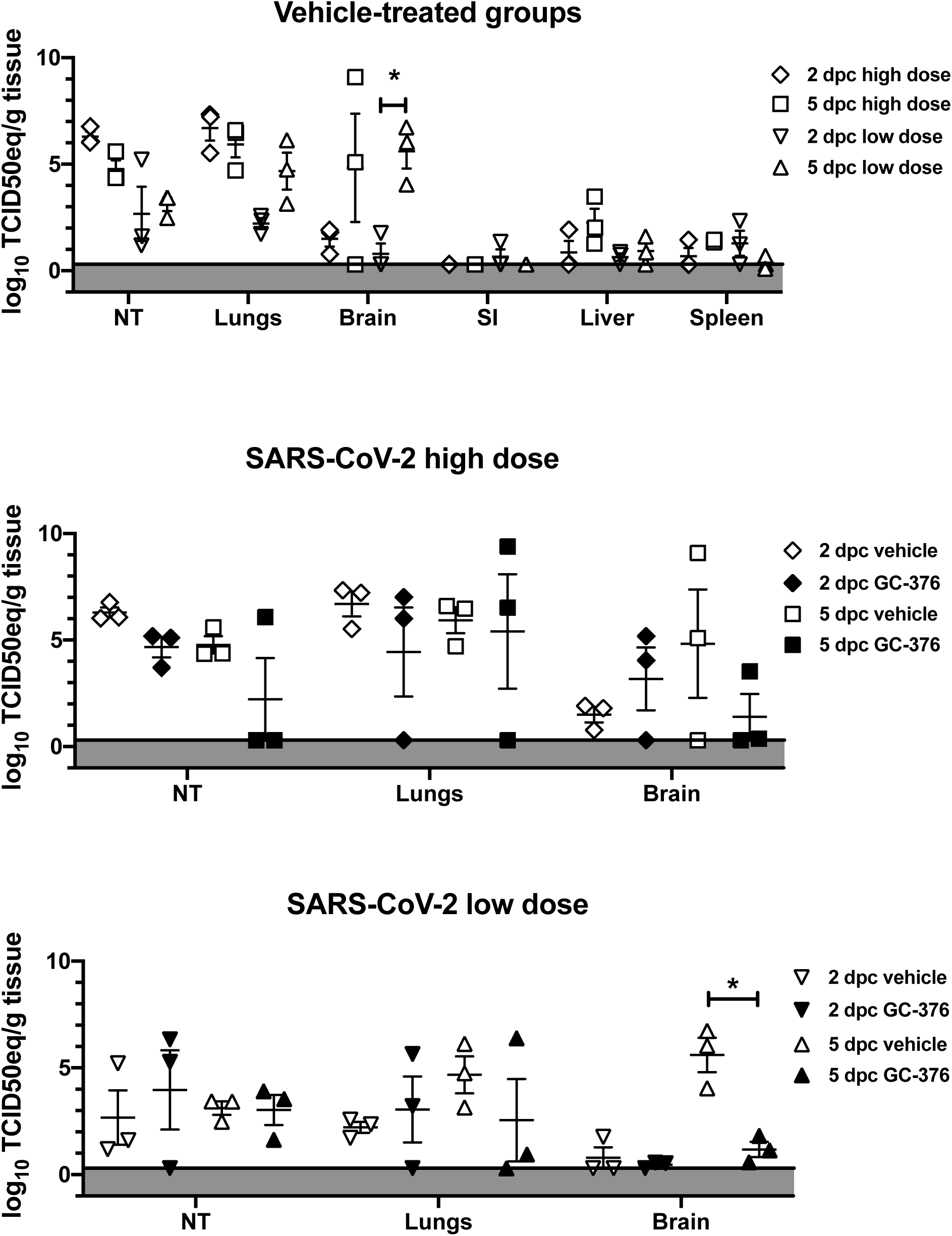
Viral loads in different mouse tissues at 2 and 5 dpc after SARS-CoV-2 challenge. Subsets of mice (n=3/group/time point) were humanely euthanized at 2 and 5 dpc. Lungs, brain, SI, liver, spleen and NT were collected. **(A)** Viral loads from G3 (high dose/vehicle) and G5 (low dose/vehicle) were determined by RT-qPCR and expressed as log_10_ TCID50 equivalent (TCID50eq) per gram of tissue. Mice challenged with high dose of SARS-CoV-2 euthanized at 2 dpc (clear diamond) and 5 dpc (clear squares) or challenged with low dose of SARS-CoV-2 euthanized at 2 dpc (clear inverse triangle) and 5 dpc (clear triangles). Viral loads in lungs, brain and NT from mice challenged with **(B)** high dose or **(C)** low dose of SARS-CoV-2 and treated with drug-vehicle (clear) or GC-376 (black) were analyzed from samples collected at 2 and 5 dpc. The mean +/-SEM is shown. Statistical analysis was performed by a two-way ANOVA with a Dunnett’s multiple comparisons test, * p<0.05

### Effects of GC-376 treatment on SARS-CoV-2-induced pathology in mice

Histopathological analysis of lungs, NT, and brains at 2, 5, and 14 dpc (Table 1) showed overall milder lesions in mice treated with GC-376 and challenged with the low dose virus (Fig. 4). At 2 dpc, no differences were observed in the two groups challenged with the high dose (G3 and G4). Lesions at 2 dpc consisted of mild-moderate interstitial lymphoplasmacytic and histiocytic inflammation especially centered around vessels and peribronchiolar spaces (Fig 4 A-B, asterisks). In addition, marked edema was effacing the alveoli and interstitium in the G3 group (red arrowheads). At 2 dpc, G5 and G6 (low dose/vehicle and low dose/GC-376) presented similar but milder extent of lymphoplasmacytic and histiocytic inflammation (Fig 4C-D, asterisks) compared to high dose groups. At 5 dpc, interstitial pneumonia in both G4 and G3 (high dose/GC-376 and high dose/vehicle) groups increased of severity and percentage of parenchyma affected (Fig. 4F-G) and again no significant differences in scoring were observed. Alveolar septal thickening and consolidation were caused by the severe proliferation of type II pneumocytes and lymphoplasmacytic inflammation (Fig. 4F-G arrowhead and asterisk, respectively). In addition, the high dose/vehicle G3 group presented a more pronounced neutrophilic inflammation (Fig 4G white arrowhead) with necrosis of alveolar septa, hemorrhage and presence of binucleated, and multinucleated epithelial cells (syncytia) sloughing into the alveolar lumen. Mild-moderate pleuritis was often observed in the affected groups. Despite the minimal differences observed at 5 dpc in the high dose challenged groups, 2 out of 3 mice treated with GC-376 and challenged with the low dose, did not present any pulmonary lesions (Fig 4H). In the low dose/vehicle group, mice presented mild to moderate lesions similar to what was already described in the previous high dose groups (G3 and G4). These also included marked alveolar edema and pleuritis (Fig 4I, red arrowhead and black arrow, respectively). Airways were mostly spared of lesions, presenting occasional blebbing and sloughing of bronchial epithelial cells into the lumen (Fig 4 F, G, and I). By 14 dpc, none to minimal lesions were observed in G4 (high dose/GC-376, only one survivor) (Fig 4K), whereas in G3 (high dose/vehicle) lesions at 14 dpc could not be evaluated because there were no survivors. At 14 dpc, the mice in G6 (low dose/GC-376) presented mild-moderate multifocal perivascular cuffing forming lymphoid nodular aggregates (Fig 4L, asterisk). The G5 low dose/vehicle mice presented sparser lymphoplasmacytic and histiocytic infiltrates similar to the one observed at day 2 dpc (Fig. 4M, asterisk).

**Table 1.**
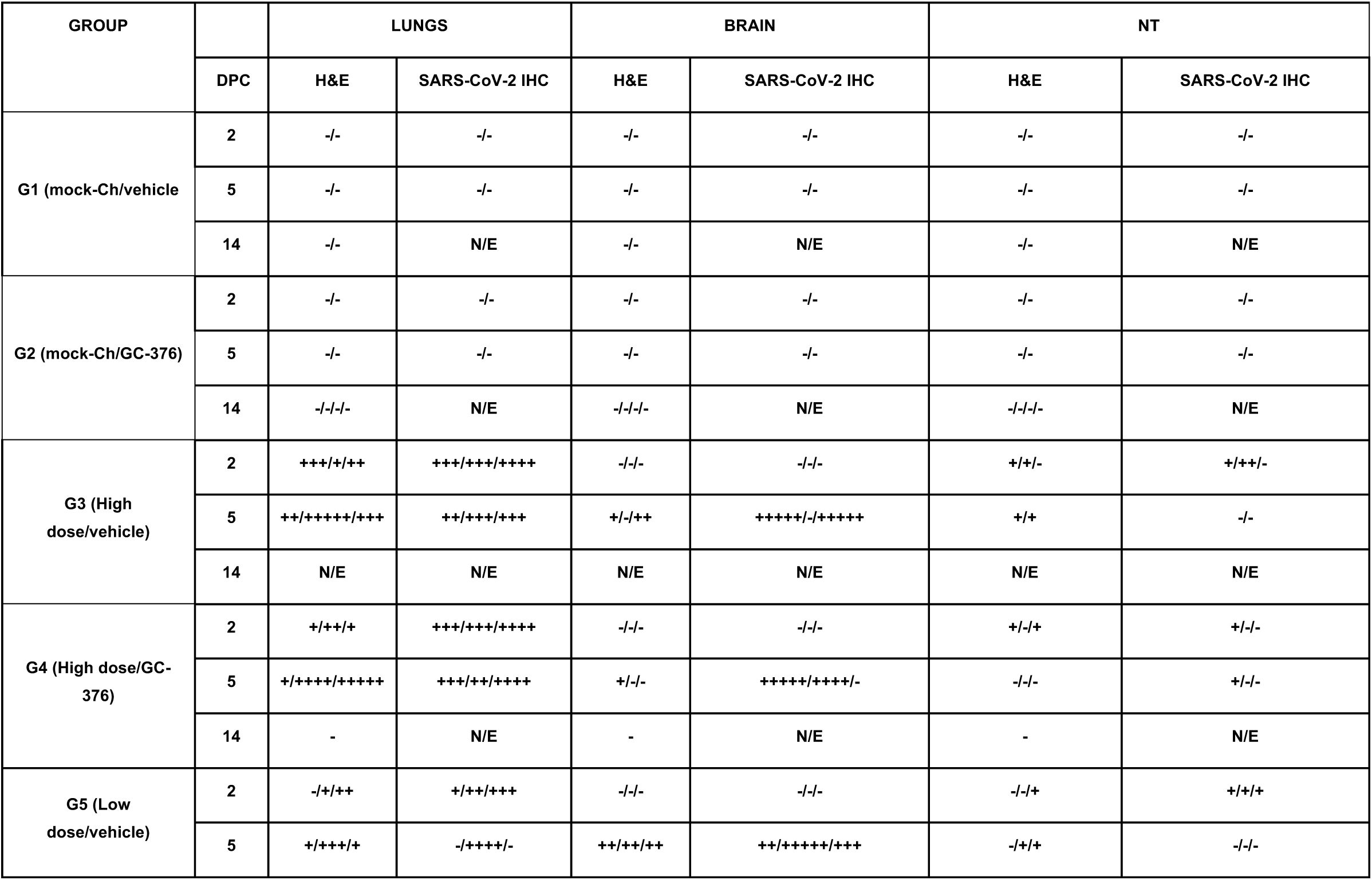
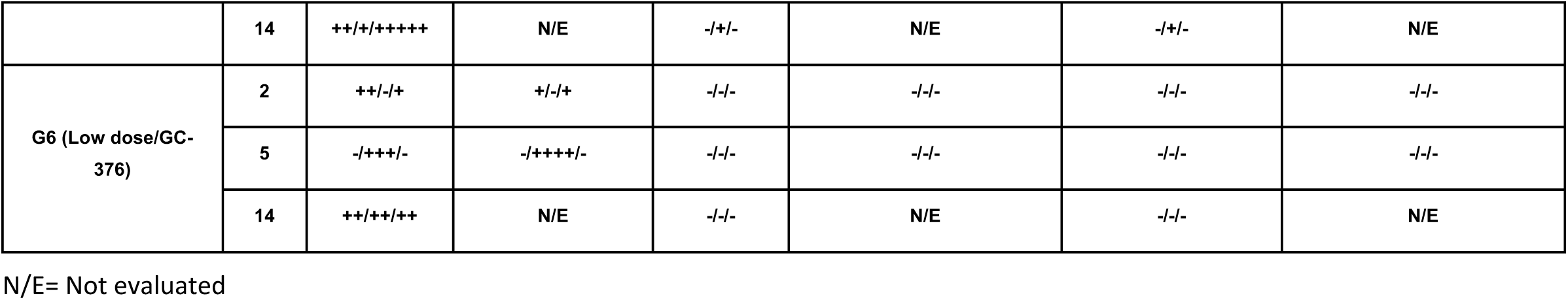
HE and SARS-CoV-2 IHC scores of the different groups evaluated in this study.

**Figure 4.**
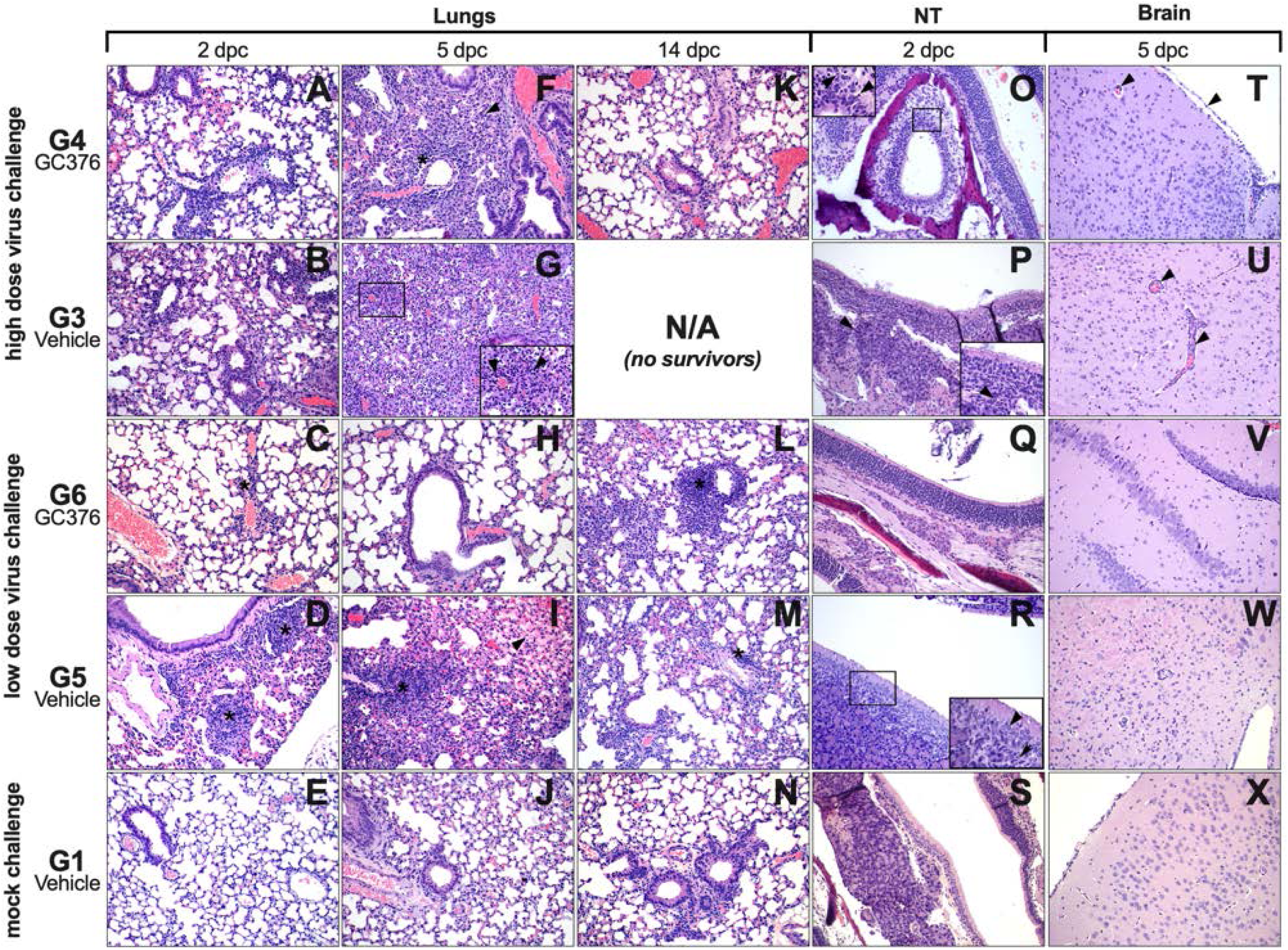
Microscopic findings in lungs, NT and brain from GC-376- and vehicle-treated mice at selected time points. **(A-D)** Mild to moderate incremental lymphoplasmacytic and histiocytic perivascular infiltrates were observed at 2 dpc in the lungs of all groups (asterisks). **(B)** G3, lungs at 2 dpc, alveolar spaces were also effaced by edema (red arrowheads). **(E)** G1, lungs at 2 dpc were normal. **(F, G and I)** Marked pulmonary consolidation was observed at 5 dpc in lungs from G4, G3 and G5 due to pneumocyte type II hyperplasia and marked interstitial inflammation (black arrowheads). Asterisks mark perivascular cuffs of mononuclear cells. **(G)** Marked neutrophilic inflammation was also observed (white arrowheads) in G3 lungs at 5 dpc **(I)** Moderate alveolar edema (red arrowhead) and pleuritis (black arrow), were present at 5 dpc in G5. **(H)** G6 lungs did not present any lesions in 2/3 mice at 5 dpc. **(J)** G1, normal lungs, 5 dpc. **(K-N)** Lungs at 14 dpc, presented from none to minimal perivascular and peri-bronchial lymphoplasmacytic and histiocytic infiltrates (asterisks). **(O, P, and R)** Nasal turbinates presented minimal changes in G4, G3 and G5 at 2 dpc. **(O)** G4, NT, at 2 dpc; mild vacuolar degeneration, pyknosis, and karyorrhexis of single olfactory epithelial cells (arrowheads). **(P)** G3, NT, at 2 dpc; the lamina propria was mildly expanded by neutrophils, lymphocytes and macrophages (arrowheads). **(Q, S)** No lesions were observed in neither G6 nor G1 NT at 2 dpc. **(R)** G5, NT at 2 dpc; olfactory epithelium was attenuated and vacuolated (arrowhead), and the lamina propria expanded by small numbers of neutrophils and macrophages (black arrow). **(T-U, W)** Brain from G4, G3 and G5 mice at 5 dpc, presented mild to moderate multifocal lymphoplasmacytic perivascular cuffing (arrowheads) surrounded by mild areas of hypercellularity (gliosis). **(V, X**) No lesions were observed in G6 and G1, at 5 dpc. Pictures and inserts were taken at 20X and at 40X magnification, respectively.

Lesions in the olfactory epithelium of the NT were minimal, only observed in 2 out of 3 mice at 2 dpc, in G4 (high dose/GC-376), G3 and G5 (high virus dose/vehicle and low virus dose/vehicle, respectively) at all timepoints, and none in G6 (low dose/GC367). These consisted of vacuolar degeneration, pyknosis and karyorrhexis of random single mucosal epithelial cells (Fig 4 O arrowheads).

In G3 (high dose/vehicle) NT lesions were accompanied by infiltration of mild numbers of neutrophils, macrophages, and lymphocytes (Fig. 4P, arrowhead). No lesions were observed in any of the mice from G6 (low dose/GC-376) (Fig 4Q) similar to G1 (Fig 4S). In contrast, G5 (low dose/vehicle) presented lesions similar to G3 (high dose/vehicle), with mild degeneration of the olfactory epithelial cells and mild mixed inflammatory infiltrate (Fig. 4R, arrow and arrowheads). At 5 dpc, the brain of mice in G4 and G3 (high dose/GC-376 and high dose/vehicle), presented similarly from none to mild microscopic lesions that consisted mostly of mild gliosis, lymphoplasmacytic and histiocytic meningoencephalitis (Fig 4, T-U arrowheads). Interestingly, none of the mice in the G6 (low dose/GC-376) group presented brain lesions (Fig 4V), as opposed to mice in G5 (low dose/vehicle) which presented lesions similar to the high dose challenge groups (Fig. 4W).

Viral antigen in lungs was readily detected in lungs of G3 and G4 (high dose/vehicle and high dose/GC376 respectively) with little variations in the distribution, prevalence in the section and intensity of the staining (Fig 5, A-B). In the case of G6 and G5 (low dose/GC-376 and low dose/vehicle), viral antigen was present, but lower in G6 compared to G5 (Fig.5 C-D). By 5 dpc, amount of viral antigen increased with no appreciable difference between high dose or low dose challenge groups (Fig 5 F-I). In lungs, viral antigen staining was observed within the cytoplasm of pneumocytes type I, II, and alveolar macrophages.

**Figure 5.**
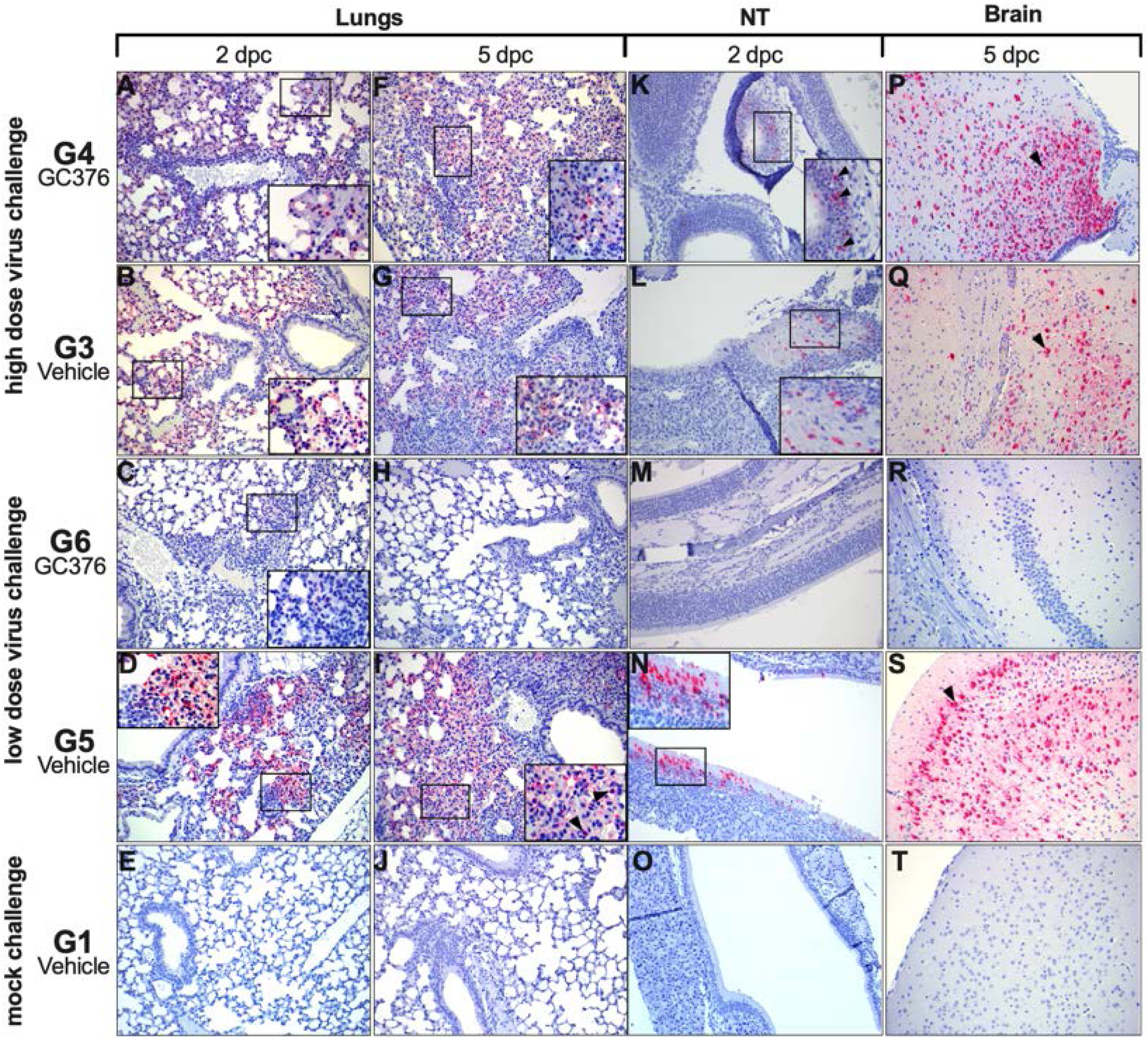
SARS-CoV-2 virus nucleocapsid tissue distribution in lungs, NT and brain analysis through IHC. **(A, F)** G4, lungs, 2 and 5 dpc, respectively; moderate amount of virus antigen (red staining) is present within pneumocyte type I, II, and alveolar macrophages. **(B, G)** G3, lungs, 2 and 5 dpc, respectively; moderate amount of virus N antigen (red staining) is present within pneumocyte type I, II. **(C, H)** G6, lungs, 2 and 5 dpc, respectively; rare pneumocytes type I and II present intracytoplasmic viral nucleocapsid at 2 dpc and normal lungs at 5 dpc. **(D, I)** G5, lungs, 2 and 5 dpc, respectively; mild numbers of cells (pneumocyte type I, II, and alveolar macrophages) present virus antigen positivity within the cytoplasm (arrowheads). **(E, J)** G1, normal lungs at 2 and 5 dpc, respectively. **(K, L)** G4 and G3, respectively, NT, 2 dpc; mild amount of intracytoplasmic virus antigen within olfactory epithelium. **(M)** G6, normal NT, 2 dpc. **(N)** G5, NT, 2 dpc; moderate amount of virus nucleocapsid present within both respiratory epithelium and olfactory epithelium. **(O)** G1, normal NT, 2 dpc **(P, Q)** G4 and G3, respectively, brain, 5 dpc; moderate to large amount of virus antigen within neurons (arrowheads) **(R)** G6, normal brain, 5 dpc. **(S)** G5, brain 5 dpc; moderate to large amount of virus antigen within neurons (arrowheads). **(T)** G1, normal brain, 5 dpc. Pictures were taken at 20X magnification and inserts were taken at 40X magnification.

In NT, viral antigen at 2 dpc was observed in 1 out of 3 mice in G4 (high dose/GC376) and in 2 out of 3 mice in G3 high dose/vehicle (Fig 5 K-L). In mice challenged with the low virus dose, viral antigen was not detected in G6 mice (low dose/GC376) whereas 3 out of 3 mice showed mild positive staining in G5 (low dose/vehicle) at 2 dpc (Fig 5 M-N). Viral antigen in NT was located in the cytoplasm of the ciliated epithelial cells of the respiratory mucosa and in the olfactory cells of the pseudostratified olfactory epithelium.

Despite the mild histopathological lesions, the brain presented extensive viral antigen positivity in neurons of G4 and G3 (high dose/GC376 and high dose/vehicle, respectively) at 5 dpc with no relevant differences (Fig. 5 P-Q, arrowheads). Viral antigen was distributed in neurons of both the white and grey matters, and randomly across the olfactory bulb, cerebrum, limbic system (e.g., hippocampus, hypothalamus, thalamus) and brainstem. Interestingly, when challenged with the low dose, no antigen was detected in any of the mice in G6 (low virus dose/GC-376) in contrast with all the mice in G5 (low virus dose/vehicle) which presented moderate to large amounts of viral antigen in the brain at 5 dpc (Fig 5 R-S). This suggested a lower SARS-CoV-2 replication preventing the spreading of the virus in the presence of GC-376. These observations are consistent with the vRNA load data by RT-qPCR for both of these groups (Fig 3C).

IHC using anti-CD3 or anti-Iba-1 antibodies was performed to better characterize the type and extent of the cellular inflammatory infiltrate, T lymphocytes and cells of the monocyte/macrophage lineage, respectively (Table 2). No significant differences in numbers of CD3+ cells in any organ were noticed between vehicle-treated mice or GC-376 treated mice challenged with the high dose SARS-CoV-2 at the given timepoints (Fig 6, panels 1-2, 6-7, 11-12, 16-17). On the contrary, a clearly lower number of CD3+ cells were observed at both 2 and 5 dpc in the lungs, NT and brain of G6 mice (low virus dose/GC-376) compared to G5 mice (low virus dose/vehicle) (Fig. 6, panels 3-4, 8-9, 13-14, 18-19). This suggests a potential implication of GC-376 in preventing more severe inflammation and virus-induced tissue damage (Fig 6; Table 2). Similar results were observed with Iba-1 IHC, showing similar staining patterns in the organs from the high dose challenge groups but lower staining overall in G6 (low dose/GC-376) when compared against its control G5 (low dose/vehicle) (Fig 6, panels 21-40). Findings were overall consistent with the extent of inflammatory reaction and lesions observed by HE.

**Table 2.**
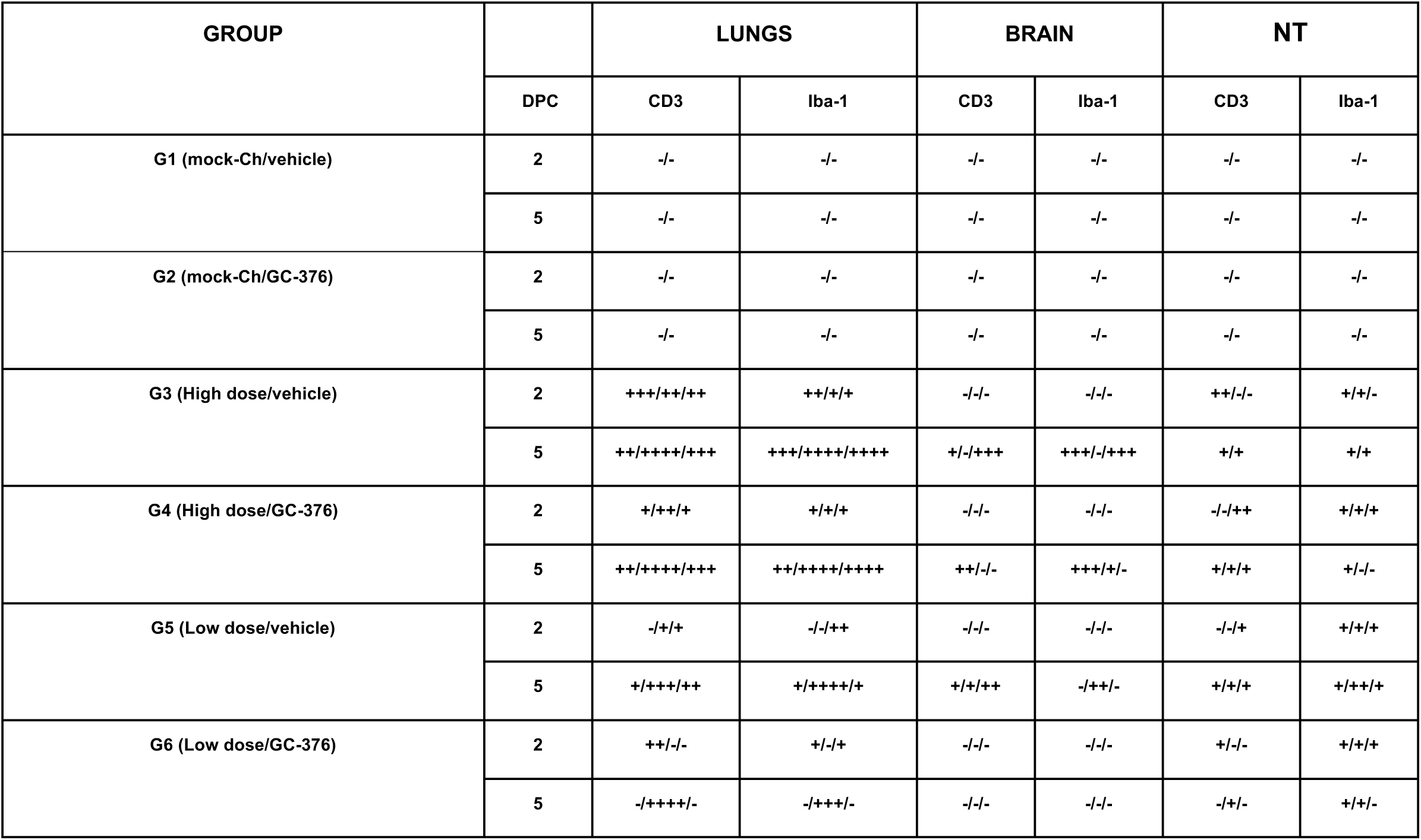
CD3 and Iba1 IHC scores of the different groups evaluated in this study.

**Figure 6.**
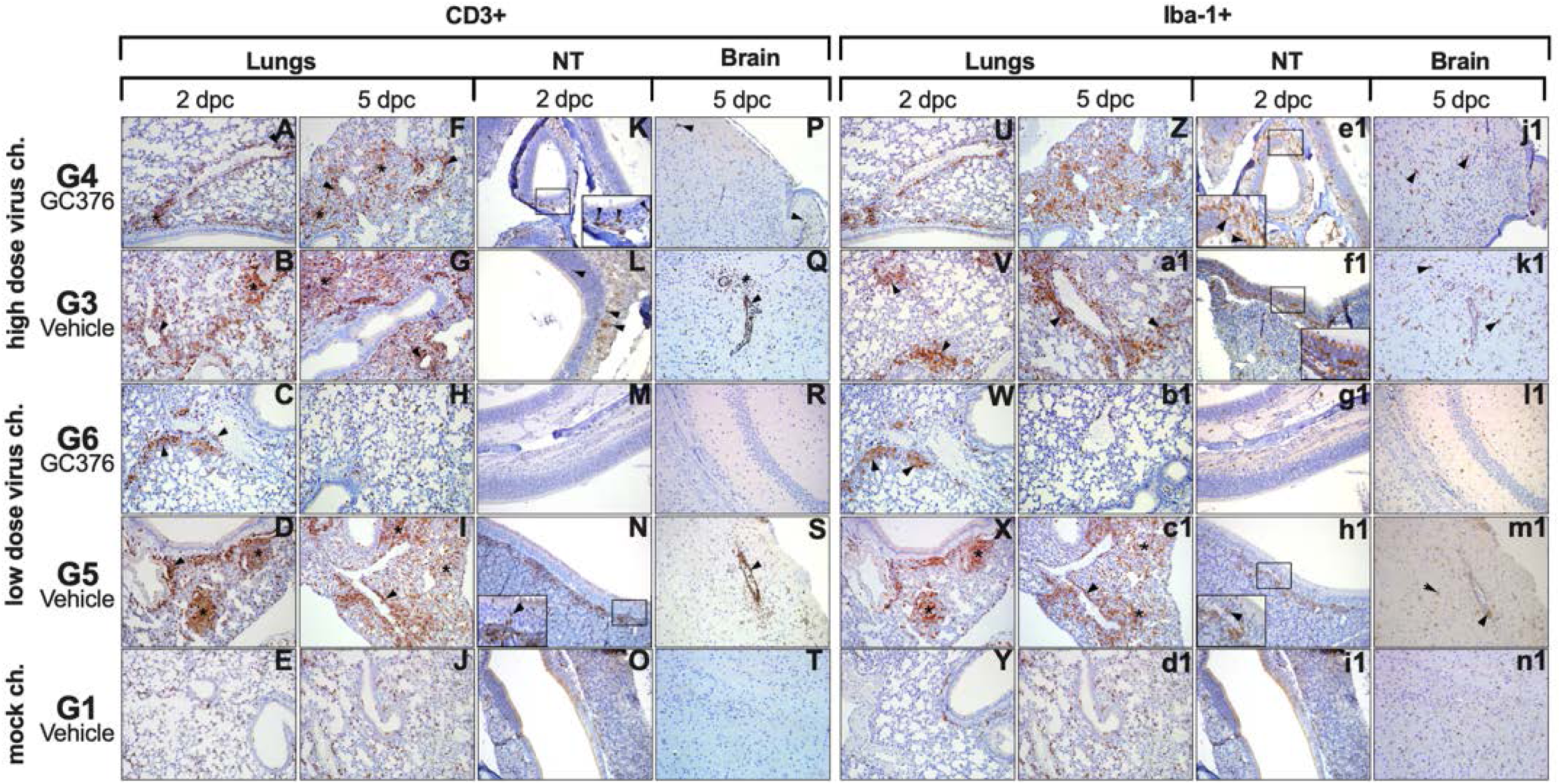
T-cell and monocyte/macrophages accumulation in lung, NT and brain from GC-376- and vehicle-treated mice at selected timepoints. **(1-4, 6-7, 9)** Increased numbers of T lymphocytes are expanding perivascular and peribronchiolar spaces (arrowheads) and invading the interstitium (asterisks) at 2 and 5 dpc in G4, G3, G6 and G5. In G6, 2/3 mice did not present any increase in T-cells at 5 dpc. **(5, 10)** G1, normal lungs from uninfected control mice at 2 and 5 dpc, respectively. Note resident CD3+ lymphocyte concentrations in the resting tissue. **(11-12, 14)** Mild increase in T-cells were observed in G4, G3 and G5 NT at 2 dpc invading lamina propria and olfactory epithelium (arrowheads). **(13, 15)** No increase in CD3+ staining was observed in G6 and G1 NT at 2 dpc. **(16-17, 19)** Mild-moderate increase in perivascular T-cells was observed in G4, G3, and G5 brains at 5 dpc. **(18, 20)** No increase in CD3+ staining was observed in G6 and G1 brains at 5 dpc. **(21-24, 26-27, 29)** Increased numbers of Iba-1+ cells were found effacing perivascular and peribronchiolar spaces (arrowheads) and invading the interstitium (asterisks) at 2 and 5 dpc in G4, G3, G6 and G5. **(28)** G6, 2/3 mice did not present any increase in T-cells at 5 dpc. **(25, 30)** G1, normal lungs from uninfected control mice at 2 and 5 dpc, respectively. Note resident Iba-1+ macrophages concentrations in the resting tissue. **(31-32, 34)** Mild increase in macrophages were observed in G4, G3 and G5 NT at 2 dpc invading lamina propria and olfactory epithelium (arrowheads and inserts). **(33, 35)** No increase in Iba-1+ staining was observed in G6 and G1 NT at 2 dpc. **(36-37, 39)** Mild-moderate increase in perivascular macrophages was observed in G4, G3, and G5 brains at 5 dpc accompanied by marked increased in staining of resident microglial cells within areas of intense virus antigen positivity (arrowheads). **(38, 40)** No increase in Iba-1+ staining was observed in G6 and G1 brains at 5 dpc. Pictures and inserts were taken at 20X and 40X magnification, respectively.

Taken together, these results suggest that GC-376 is able to prevent to some extent the inflammatory reactions induced by SARS-CoV-2 infection.

## DISCUSSION

In the last two decades, coronaviruses have been implicated in significant zoonotic outbreaks (SARS-CoV-1 in 2003 and MERS in 2012) and in the COVID-19 pandemic (SARS-CoV-2 in 2019/2020). The current COVID-19 pandemic has caused more than 45 million confirmed infections associated to more than 1 million deaths worldwide, a public health emergency of incomparable proportions in the 21^st^ century ^1^. Many anti-SARS-CoV-2 antiviral efforts have focused on the repurposing of antivirals compounds rather than the development of novel compounds. Among those, drugs such as hydroxychloroquine (HQC), remdesivir, favipinavir, ivermectin, among others have been proposed as promising candidates. There are currently more than 50 different clinical trials where compounds with potential antiviral activity are under evaluation ^26^. Previous studies have shown HQC antiviral activity *in vitro* against SARS-CoV-1 and, early during the COVID-19 pandemic, against SARS-CoV-2 ^27,28^. However, further analysis in animal models (hamsters and monkeys) and in human clinical trials showed that HCQ is ineffective against SARS-CoV-2 in vivo ^29-31^. More promising outcomes have been observed with remdesivir, an adenosine nucleoside triphosphate analog, that shows antiviral activity *in vitro* and in mice against a surrogate chimeric SARS-CoV-2 virus ^13,32-34^. More importantly, remdesivir was shown to shorten the time to recovery of COVID-19 hospitalized patients ^35,36^. The FDA has recently approved the remdesivir for treatment of COVID-19 patients, ages ≥ 12 years old and weighing ≥40 pounds, that require hospitalization. However, remdesivir may offer limited benefits to those that are severely ill. More importantly, a press release from the WHO on November 20, 2020 recommends against the use of remdesivir in COVID-19 patients. Immunotherapy using convalescent plasma or monoclonal antibodies is another avenue of great focus in the treatment of COVID-19. In this regard, the FDA has recently released an emergency use authorization for the investigational monoclonal antibody therapy Bamlanivimab (LY-CoV55, Elli Lilly) to treat mild-to-moderate COVID-19 in adult and pediatric patients with high risk to become severely ill or hospitalized (typically those ≥65 years of age or who have certain chronic medical conditions). It is important to mention that neither bamlanivimab nor REGN-COV2 (Regeneron) immunotherapies have proven effective in COVID-19 patients requiring mechanical ventilation (^37,38^ and NCT04426695). Thus, there is still a long road ahead in the development of adequate countermeasures against the SARS-CoV-2 virus.

Herein we evaluated the antiviral activity *in vivo* of GC-376, a compound that has shown antiviral activity against SARS-CoV-2 *in vitro*. For these studies we used the K18-hACE2 mouse model which was previously established as a suitable small animal model of SARS-CoV-1 infection ^18^. We must note that at the time we initiated our studies, no reports were available in the public domain regarding the suitability of the K18-hACE2 mouse model to study SARS-CoV-2 infection. Since then, several reports have emerged and our results are in agreement with the notion that the K18-hACE2 mouse model is a suitable small animal model to study SARS-CoV-2 pathogenesis _^20,21,25^_. Previous reports have also shown that K18-hACE2 mice are amenable for the study of vaccines against SARS-CoV-2 ^24^ but, to our knowledge, assessment of this mouse model for the evaluation of potential antivirals against SARS-CoV-2 was lacking. We observed clinical signs of disease and mortality in K18-hACE2 mice upon SARS-CoV-2 challenge in a virus dose-dependent manner (Figs 3 and 4). Severe clinical signs and 0% survival was observed at the high virus challenge dose in contrast to the relatively delayed onset of clinical signs and 60% survival at the low virus challenge dose demonstrating that this model is not hypersusceptible to the SARS-CoV-2 virus. Data on clinical signs and body weight monitoring indicated that mice in the low virus challenge dose were indeed infected and those that survived were essentially devoid of clinical signs by 14 dpc. This is relevant since the K18hACE2/SARS-CoV-2 model offers the opportunity to study disease outcome and host responses which in humans are thought to be related, at least in part, to the amount of virus during exposure. In contrast, other mouse models, either transduced with a recombinant adenovirus or modified with CRISP/Cas9 to express hACE2 have shown limited susceptibility to SARS-CoV-2 and/or neither apparent clinical signs nor mortality ^2,39^.

The toxicity of GC-376 (7-day course at 40 mg/kg/day) was evaluated through changes in body weight and survival (Fig 2) and histopathological changes in different tissues at 7 days post-treatment (14 dpc). These analyses indicated lack of obvious acute toxicity of GC-376 in K18-hACE-2 mice. Although previous studies in cats showed no GC-376 associated toxicity those were performed using lower doses than in our studies. And in studies using GC-376 against murine hepatitis virus (MHV) A59 in Balb/c mice no specific toxicity evaluations were performed ^6,7,23^.

We observed a modest although clearly beneficial effect of GC-376 against SARS-CoV-2 *in vivo* based on the collective analysis of the following observations: Slight delay in onset of clinical signs accompanied by one survivor in the GC-376-treated group compared to no survivors in the vehicle-treated mice challenged with a high virus dose. More interindividual variations with several more tissue samples showing vRNA below levels of detection comparing GC-376-treated to vehicle-treated animals challenged with either high or low virus dose. Although not statistically significant, average lower vRNA levels were observed in tissue samples from the NT, lungs, and brain in mice treated GC-376 than in the vehicle-treated counterparts. Faster body weight recovery and significantly reduced vRNA presence in brain samples collected at 5 dpc was observed in GC-376-treated versus vehicle-treated mice challenged with low dose virus. The reduced vRNA load in brain samples was consistent with reduced viral antigen staining. Reduced virus involvement in the brain is important considering the neurologic features observed in COVID-19 patients where disorientation, poorly organized movements and confusion among other neurological symptoms have been observed ^40^. Additionally, COVID-19 neurological symptoms can be potentially life threatening in around 65% of the cases ^41^. A positive effect of GC-376 was also supported by the histopathological examinations and analysis of markers of cellular infiltration (CD3 and Iba-1). Milder lesions and lower detection of CD3^+^ and Iba-1^+^ cells in lungs, NT and brain samples were observed in GC-376-treated mice compared to vehicle-treated mice. The decrease in CD3^+^ and Iba-1^+^ cells was correlated with milder inflammation also observed at HE. A lower, measured inflammatory response is thought to be favorable for the host, in contrast with an excessive inflammatory reaction, such as the multisystemic inflammatory syndrome, which is considered a major complication of SARS-CoV-2 infection, often fatal in COVID-19 patients. The widespread virus colonization of neurons and associated meningoencephalitis observed in mice, partially explains the neurological deficits observed COVID-19 patients ^41,42^. Various infective mechanisms and paths can be exploited by the virus to reach the central nervous system. SARS-CoV-2 can access the brain through various neural or hematogenous routes that connect nasal mucosae, primary site of virus replication, to the anatomically adjacent forebrain across the cribriform plate of the ethmoid bone ^43^. Our results are in agreement with the notion that virus dissemination to the brain is likely to occur from the upper respiratory tract via non-neuronal cells that provide metabolic and structural support to sensory olfactory nerve cells ^44^. This explains why anosmia (loss of smell) is one of the hallmark symptoms of COVID-19 ^44^.

The modest *in vivo* antiviral activity of GC-376 in SARS-CoV-2-infected K18hACE2 mice is consisted with its moderate *in vitro* antiviral activity. The reported *in vitro* EC_50_ values of GC-376 against SARS-CoV-2 in cell culture range from 0.5 to 3.4 µM from several independent studies, depending on the cell types and assay conditions^12,13,45^. In comparison, a GC-376 analog **6j** was recently reported to improve survival in MERS-CoV-infected mice, and the EC_50_ value of the *in vitro* cellular antiviral activity of **6j** against MERS-CoV was 0.04 ± 0.02 µM^10^. This result suggests that the *in vitro* cellular antiviral activity of GC-376 against SARS-CoV-2 needs to be improved by 10–100-fold to achieve desired *in vivo* antiviral efficacy. As such, the lead optimization on GC-376 should mainly focus on improving the cellular antiviral activity. Alternatively, higher doses of GC-376, alone or in combination with other antivirals such as remdesivir or EIDD-2801 and/or anti-inflammatory drugs such as dexamethasone could be evaluated against SARS-CoV-2. Although such studies are beyond the scope of the present report, they are undoubtedly warranted. This is particularly the case since currently approved intervention strategies are limited and not very efficacious to treat severe COVID-19 cases. To our knowledge, ours is the first to study the *in vivo* efficacy of GC-376 against SARS-CoV-2. The precedent successful examples of GC-376 in treating FIP CoV infection and **6j** in treating MERS-CoV infection in mice, coupled with the promising anti-inflammatory effects of GC-376 against SARS-CoV-2 infection in mice from this current study, warrants further development of GC-376 to specifically target SARS-CoV-2.

## ACKNOWLEDGMENTS

We thank the personnel from the Animal Health Research Center and the Histology laboratory, College of Veterinary Medicine, University of Georgia. This study was supported by a subcontract from the Center for Research on Influenza Pathogenesis (CRIP) to D.R.P. under contract HHSN272201400008C from the National Institute of Allergy and Infectious Diseases (NIAID) Centers for Influenza Research and Surveillance (CEIRS). J. W. thanks the support from the National Institutes of Health (NIH) (Grants AI147325 and AI157046) and the Arizona Biomedical Research Centre Young Investigator grant (ADHS18-198859).

## References

1 WHO. https://www.who.int/emergencies/diseases/novel-coronavirus-2019. (2020).

2 Hassan, A. O. et al. A SARS-CoV-2 Infection Model in Mice Demonstrates Protection by Neutralizing Antibodies. Cell 182, 744–753 e744, doi:10.1016/j.cell.2020.06.011 (2020).

3 Corbett, K. S. et al. Evaluation of the mRNA-1273 Vaccine against SARS-CoV-2 in Nonhuman Primates. N Engl J Med, doi:10.1056/NEJMoa2024671 (2020).

4 Smith, T. R. F. et al. Immunogenicity of a DNA vaccine candidate for COVID-19. Nat Commun 11, 2601, doi:10.1038/s41467-020-16505-0 (2020).

5 Ahidjo, B. A., Loe, M. W. C., Ng, Y. L., Mok, C. K. & Chu, J. J. H. Current Perspective of Antiviral Strategies against COVID-19. ACS Infect Dis 6, 1624–1634, doi:10.1021/acsinfecdis.0c00236 (2020).

6 Pedersen, N. C. et al. Efficacy of a 3C-like protease inhibitor in treating various forms of acquired feline infectious peritonitis. J Feline Med Surg 20, 378–392, doi:10.1177/1098612X17729626 (2018).

7 Kim, Y. et al. Reversal of the Progression of Fatal Coronavirus Infection in Cats by a Broad-Spectrum Coronavirus Protease Inhibitor. PLoS Pathog 12, e1005531, doi:10.1371/journal.ppat.1005531 (2016).

8 Li, Q. & Kang, C. Progress in Developing Inhibitors of SARS-CoV-2 3C-Like Protease. Microorganisms 8, pdoi:10.3390/microorganisms8081250 (2020).

9 Kim, Y. et al. Broad-spectrum antivirals against 3C or 3C-like proteases of picornaviruses, noroviruses, and coronaviruses. J Virol 86, 11754–11762, doi:10.1128/JVI.01348-12 (2012).

10 Rathnayake, A. D. et al. 3C-like protease inhibitors block coronavirus replication in vitro and improve survival in MERS-CoV-infected mice. Sci Transl Med 12, pdoi:10.1126/scitranslmed.abc5332 (2020).

11 Vuong, W. et al. Feline coronavirus drug inhibits the main protease of SARS-CoV-2 and blocks virus replication. Nat Commun 11, 4282, doi:10.1038/s41467-020-18096-2 (2020).

12 Ma, C. et al. Boceprevir, GC-376, and calpain inhibitors II, XII inhibit SARS-CoV-2 viral replication by targeting the viral main protease. Cell research 30, 678–692, doi:10.1038/s41422-020-0356-z (2020).

13 Fu, L. et al. Both Boceprevir and GC376 efficaciously inhibit SARS-CoV-2 by targeting its main protease. Nat Commun 11, 4417, doi:10.1038/s41467-020-18233-x (2020).

14 Munster, V. J. et al. Respiratory disease in rhesus macaques inoculated with SARS-CoV-2. Nature, doi:10.1038/s41586-020-2324-7 (2020).

15 Sia, S. F. et al. Pathogenesis and transmission of SARS-CoV-2 in golden hamsters. Nature 583, 834–838, doi:10.1038/s41586-020-2342-5 (2020).

16 Imai, M. et al. Syrian hamsters as a small animal model for SARS-CoV-2 infection and countermeasure development. Proc Natl Acad Sci U S A 117, 16587–16595, doi:10.1073/pnas.2009799117 (2020).

17 Schlottau, K. et al. SARS-CoV-2 in fruit bats, ferrets, pigs, and chickens: an experimental transmission study. Lancet Microbe, doi:10.1016/S2666-5247(20)30089-6 (2020).

18 McCray, P. B., Jr. et al. Lethal infection of K18-hACE2 mice infected with severe acute respiratory syndrome coronavirus. J Virol 81, 813–821, doi:10.1128/JVI.02012-06 (2007).

19 Rathnasinghe, R. et al. Comparison of Transgenic and Adenovirus hACE2 Mouse Models for SARS-CoV-2 Infection. bioRxiv, doi:10.1101/2020.07.06.190066 (2020).

20 Yinda, C. K. et al. K18-hACE2 mice develop respiratory disease resembling severe COVID-19. bioRxiv, doi:10.1101/2020.08.11.246314 (2020).

21 Golden, J. W. et al. Human angiotensin-converting enzyme 2 transgenic mice infected with SARS-CoV-2 develop severe and fatal respiratory disease. JCI Insight 5, pdoi:10.1172/jci.insight.142032 (2020).

22 Reed, L. M., H. A SIMPLE METHOD OF ESTIMATING FIFTY PER CENT ENDPOINTS. American Journal of Epidemiology 27, 493–497 (1938).

23 Kim, Y. et al. Broad-spectrum inhibitors against 3C-like proteases of feline coronaviruses and feline caliciviruses. J Virol 89, 4942–4950, doi:10.1128/JVI.03688-14 (2015).

24 Seo, S. H. & Jang, Y. Cold-Adapted Live Attenuated SARS-Cov-2 Vaccine Completely Protects Human ACE2 Transgenic Mice from SARS-Cov-2 Infection. Vaccines (Basel) 8, pdoi:10.3390/vaccines8040584 (2020).

25 Oladunni, F. S. et al. Lethality of SARS-CoV-2 infection in K18 human angiotensin-converting enzyme 2 transgenic mice. Nat Commun 11, 6122, doi:10.1038/s41467-020-19891-7 (2020).

26 NIH. https://clinicaltrials.gov. (2020).

27 Liu, J. et al. Hydroxychloroquine, a less toxic derivative of chloroquine, is effective in inhibiting SARS-CoV-2 infection in vitro. Cell Discov 6, 16, doi:10.1038/s41421-020-0156-0 (2020).

28 Weston, S. et al. Broad Anti-coronavirus Activity of Food and Drug Administration-Approved Drugs against SARS-CoV-2 In Vitro and SARS-CoV In Vivo. J Virol 94, doi:10.1128/JVI.01218-20 (2020).

29 Kaptein, S. J. F. et al. Favipiravir at high doses has potent antiviral activity in SARS-CoV-2-infected hamsters, whereas hydroxychloroquine lacks activity. Proc Natl Acad Sci U S A, doi:10.1073/pnas.2014441117 (2020).

30 Maisonnasse, P. et al. Hydroxychloroquine use against SARS-CoV-2 infection in non-human primates. Nature 585, 584–587, doi:10.1038/s41586-020-2558-4 (2020).

31 Boulware, D. R. et al. A Randomized Trial of Hydroxychloroquine as Postexposure Prophylaxis for Covid-19. N Engl J Med 383, 517–525, doi:10.1056/NEJMoa2016638 (2020).

32 Wang, M. et al. Remdesivir and chloroquine effectively inhibit the recently emerged novel coronavirus (2019-nCoV) in vitro. Cell Res 30, 269–271, doi:10.1038/s41422-020-0282-0 (2020).

33 Pruijssers, A. J. et al. Remdesivir Inhibits SARS-CoV-2 in Human Lung Cells and Chimeric SARS-CoV Expressing the SARS-CoV-2 RNA Polymerase in Mice. Cell Rep 32, 107940, doi:10.1016/j.celrep.2020.107940 (2020).

34 Hashemian, S. M., Farhadi, T. & Velayati, A. A. A Review on Remdesivir: A Possible Promising Agent for the Treatment of COVID-19. Drug Des Devel Ther 14, 3215–3222, doi:10.2147/DDDT.S261154 (2020).

35 Beigel, J. H. et al. Remdesivir for the Treatment of Covid-19 - Final Report. N Engl J Med, doi:10.1056/NEJMoa2007764 (2020).

36 Hongchao Pan, P. D., Richard Peto, F.R.S., Quarraisha Abdool Karim, Ph.D., Marissa Alejandria M.D., M.Sc., Ana Maria Henao-Restrepo, M.D., M.Sc., César Hernández García M.D., Ph.D., Marie-Paule Kieny Ph.D., Reza Malekzadeh M.D., Srinivas Murthy M.D. C.M., Marie-Pierre Preziosi M.D., Ph.D., Srinath Reddy M.D., D.M., Mirta Roses Periago M. D., Vasee Sathiyamoorthy B.M.B.Ch., Ph.D., John-Arne Røttingen M.D., Ph.D., and Soumya Swaminathan M.D. Repurposed antiviral drugs for COVID-19 –interim WHO SOLIDARITY trial results. MedRxiv (2020).

37 Baum, A. et al. REGN-COV2 antibodies prevent and treat SARS-CoV-2 infection in rhesus macaques and hamsters. Science, doi:10.1126/science.abe2402 (2020).

38 Chen, P. et al. SARS-CoV-2 Neutralizing Antibody LY-CoV555 in Outpatients with Covid-19. N Engl J Med, doi:10.1056/NEJMoa2029849 (2020).

39 Sun, S. H. et al. A Mouse Model of SARS-CoV-2 Infection and Pathogenesis. Cell Host Microbe 28, 124–133 e124, doi:10.1016/j.chom.2020.05.020 (2020).

40 Helms, J. et al. Neurologic Features in Severe SARS-CoV-2 Infection. N Engl J Med 382, 2268–2270, doi:10.1056/NEJMc2008597 (2020).

41 Bertram, B. et al. Research Square, doi:10.21203/rs.3.rs-63687/v1 (2020).

42 Merad, M. & Martin, J. C. Pathological inflammation in patients with COVID-19: a key role for monocytes and macrophages. Nat Rev Immunol 20, 355–362, doi:10.1038/s41577-020-0331-4 (2020).

43 Barrantes, F. J. Central Nervous System Targets and Routes for SARS-CoV-2: Current Views and New Hypotheses. ACS Chem Neurosci 11, 2793–2803, doi:10.1021/acschemneuro.0c00434 (2020).

44 Brann, D. H. et al. Non-neuronal expression of SARS-CoV-2 entry genes in the olfactory system suggests mechanisms underlying COVID-19-associated anosmia. Sci Adv 6, pdoi:10.1126/sciadv.abc5801 (2020).

45 Sacco, M. D. et al. Structure and inhibition of the SARS-CoV-2 main protease reveals strategy for developing dual inhibitors against M(pro) and cathepsin L. Sci Adv, doi:10.1126/sciadv.abe0751 (2020).

